# Optimizing fungicide deployment in a connected crop landscape while balancing epidemic control and environmental sustainability

**DOI:** 10.1101/2024.12.16.628632

**Authors:** Andrea Radici, Davide Martinetti, Daniele Bevacqua

## Abstract

Bioaggressors cause significant losses in crop production, and the efficacy of control methods, primarily based on chemical compounds, comes with considerable environmental and health costs. Plant protection practices implemented locally undermine the mobility of bioaggressors, which can spread between fields, connecting different crop populations. As a consequence, the yield in a given field depends also on the management of connected fields. In this study, we evaluate the efficiency of different fungicide deployment strategies across a national-scale agriculture landscape, balancing the conflicting objectives of maximizing crop production and reducing fungicide use. We use a climate-driven metapopulation model describing the dynamics of the peach (*Prunus persica*)-brown rot (caused by *Monilinia* spp.) pathosystem in continental France. Fungicide deployment strategies are based on indices or algorithms, considering network topology, epidemic risk, territory, and stochastic sampling, which prioritize sites to be treated first. Eventually, we focus solely on the objective of maximizing harvest revenue, assuming that untreated fruit can be marketed at higher prices. The optimal strategy depends on the treatment allocation threshold: if up 20% of the area is treated, epidemic risk provides the most effective prioritization. If more than 40% of the area can be treated, a combination of random sampling and risk-based prioritization proves optimal. When considering a single objective, we find that the higher the consumer’s willingness to pay for untreated fruit, the larger the proportion of untreated sites becomes. Fungicide use could be avoided if untreated fruit were sold at 2.9 times the price of treated fruit.

## 1 Introduction

A critical obstacle to stable and reliable food systems is the threat posed by pests and pathogens (including bacteria, fungi, and insects - hereinafter referred to as “bioaggressors”), which endanger crop health (Ristaino et al., 2021). They cause production losses ranging from 17% to 30% in major crops, such as wheat, rice, maize, potatoes, and soybeans (Savary et al., 2019). Conventional bioaggressor control primarily relies on phytosanitary products (Sumberg and Giller, 2022), synthetic compounds designed to eliminate or inhibit bioaggressor growth (EPPO, 2004). While these chemicals can enhance short-term yields, they often lead to resistance in target bioaggressors (Savary et al., 2019), and their persistence in the environment poses risks to ecosystems and human health (Rosic et al., 2020; Yadav and Devi, 2017).

The spread of bioaggressors is influenced by spatial factors, including habitat size and connectivity (Rusch et al., 2010). Movement can occur via human activity (*e.g.*, trade; Hernández Nopsa et al., 2015), natural vectors (*e.g.*, insects; Strona et al., 2017) or abiotic factors (*e.g.*, wind; Meyer et al., 2017). It is worth noting that agricultural landscapes can be modeled as networks where fields (nodes) are interconnected by bioaggressor movement (edges; Gilligan, 2008; Radici et al., 2023a). This network-based approach enhances our understanding of, and ability to optimize, control strategies at multiple scales. A substantial body of literature supports the use of network models to optimize epidemiological control for animal and human diseases (Keeling and Rohani, 2011). Although scarcer, there exist applications to plant bioaggressor control. For instance, Strona et al. (2017) found that removing nodes with the highest PageRank (a measure of importance of a node based on the number and quality of edges connected to it; Page, 1998) reduces the size of the the largest set of connected nodes faster that using other methods, thus reducing the spread of *X. fastidiosa* among olive orchards in southern Italy. In context of seed markets, Andersen et al. (2019) suggested that nodes characterized by high degree (*i.e.*, which are more connected) may be identified as influential spreaders, and so prioritized to be immunized.

The objective of this study is to provide optimal management strategies to control the spread of airborne diseases. With optimal, we intend strategies seeking to maximize crop production while reducing appeal to phytosanitary products. We use a climate-driven metapopulation model that captures the spatio-temporal dynamics of a fungal disease (Radici et al., 2024). As study case, we use the peach (*Prunus persica*)-brown rot (caused by *Monilinia* spp.) pathosystem at the national scale in France. Our model subdivides French peach-growing regions into spatial cells, simulating disease dynamics influenced by local weather conditions that affect peach phenology, pathogen etiology, and the global wind-driven dispersal of *Monilinia* spores. For this pathosystem, we derive prioritization strategies for fungicide application, identifying which cells to treat first based on network topology, risk and territorial indices, and stochastic sampling. We compare the effectiveness of these strategies in reducing fungicide use while meeting production targets. Finally, we investigate the impact of different pricing scenarios for treated versus untreated fruit, estimating the price increase needed for untreated fruit to eliminate fungicide application.

## 2 Materials and methods

### 2.1 Metapopulation model overview

We used the metapopulation model presented in Radici et al. (2024) to simulate brown rot spread in peach cultivated fields in France. In the following we report an overview of the model, details can be found in the original article.

The geographic domain corresponds to the Safran grid (Bertuzzi and Clastre, 2022), consisting of square cells measuring 0.11^◦^ × 0.11^◦^ (approximately 8 × 8 km², hereinafter referred to as “nodes”), overlaid on continental France. We focused on the 755 nodes with significant peach orchard coverage (greater than 0.01 ha/km²; Fig. 1a). For each node and each year between 1996 and 2020, we calculated the ripening period, from pit hardening (*t*_0_) to harvest (*t_H_*), using a phenological temperature-dependent model (see Vanalli et al. 2021 for details). This period corresponds to the time when fruits are susceptible to infection. Note that *t_H_* varies across peach cultivars (*e.g.*, early, mid-early, mid-late, and late). For each node *i*, we ran a climate driven Susceptible-Exposed-Infected (SEI type) epidemiological model (see Bevacqua et al., 2023 for details) from *t*_0_*_,i_* to *t_H,i_*, where *I*(*t*_0_*_,i_*) = 0 *fruit/m*^2^ and *S*(*t*_0_*_,i_*) + *E*(*t*_0_*_,i_*) = 15 *fruit/m*^2^. Each year, for each node, we stochastically determined the value of *E_i_*(*t*_0_) based on the disease incidence in the previous year. If *E_i_*(*t*_0_) *>* 0 *fruit/m*^2^, the node is considered “exposed”, and the epidemic dynamics operate independently of the epidemic state of other nodes. On the other hand, if *E_i_*(*t*_0_) = 0 *fruit/m*^2^, no epidemic occurs until an inoculum from connected infected nodes is introduced. In this case, we calculate the daily probability of *E_i_*(*t*) becoming positive (*E_i_*(*t*) *>* 0 *fruit/m*^2^). Such probability depends on the epidemic status of the other nodes (*i.e.*, higher infection levels translate into higher spore production) and on the airborne connectivity expressed via a time-varying connectivity matrix **W_t_**, where the element *w_ijt_* represents the probability that spores released from node *i* are deposited in node *j* on day *t*. The connectivity matrix has been computed multiple averaging Lagrangian trajectory simulations performed with HYSPLIT (Draxler and Hess, 1998), integrated with an aerobiological model simulating the transport of *Monilinia* spores, accounting for environmental conditions for spore advection, survival and deposition. In Fig. 1b we display a simplified version of **W_t_** into its static equivalent **W** by averaging over time during the ripening period. The matrix **W** summarizes the connectivity of a network where nodes represent spatial cells, and wind-driven spore transport forms the directed and weighted edges connecting these nodes.

**Figure 1:**
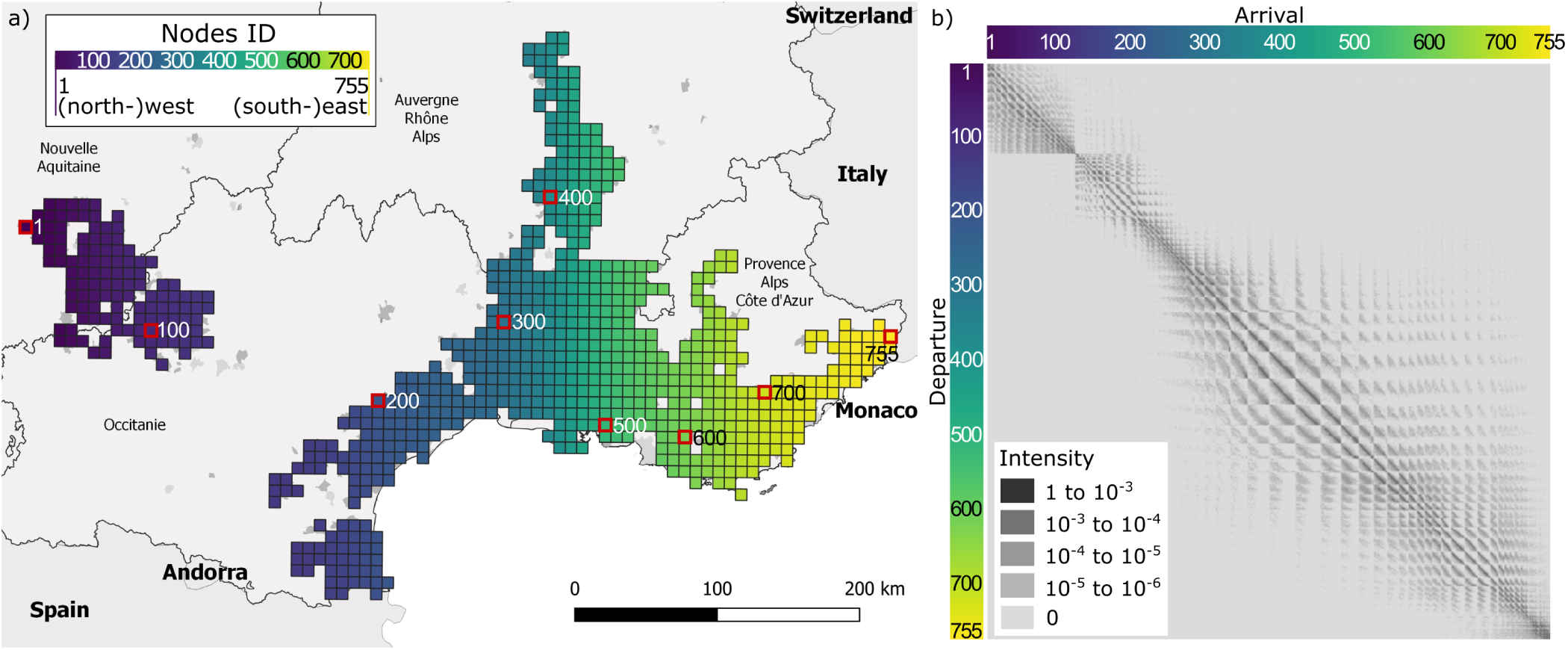
a) The study domain, consisting of 755 square cells (nodes) overlaid on the peach production basin in France, and b) the corresponding wind-connectivity matrix **W**. The node identifiers (ID) range from 1 to 755, with some reference sites indicated (IDs 1, 100, 200, 300, 400, 500, 600, 700, and 755). The IDs increase first from west to east, and second, from north to south.

### 2.2 Strategies to prioritize nodes to be treated

A disease control action, such as the application of fungicide, is associate with costs. If these costs cannot be fully covered due to economic, environmental, or social reasons, it is essential to identify the nodes where immunization should be prioritized. We therefore define a “strategy” as a ranking of nodes, obtained via an index or an algorithm, which prioritize nodes to be treated first. Consequently, we define an “index” as the quantification of a property of a node, which univocally defines its ranking (*e.g.*, by decreasing order of the index), and an “algorithm” as a procedure to sample nodes within the network. In this work, we concentrate on stochastic algorithms only.

There are several indices and algorithms that have been proposed to rank the nodes of a network for targeted interventions aimed at preventing undesirable spreading phenomena, such as diseases among organisms, malware across computers, or rumors between people (de Arruda et al., 2014). In the case of a network of orchards, addressing a node *i* involves preventing the spread of the disease within *i* by treating its cultivated area with fungicide (*E_i_*(*t*) = *I_i_*(*t*) = 0 *fruit/m*^2^).

We propose 14 different control strategies based on *i*) network centrality indices, such as in-degree, in-strength, out-degree, out-strength, and betweenness; *ii*) network propagation indices, such as coreness, GRWA, voteRank; *iii*), epidemiologial risk indices, *i.e.* vulnerability and dangerousness, (see Radici et al., 2024, for details); *iv*) host density indices, *v*) random and regular sampling, and *vi*) mixing previous criteria (Sutrave et al., 2012). Namely, we derived the vulnRand and the danRand algorithms where sampling probability is weighted by vulnerability and dangerousness, respectively. The list of all considered strategies with a brief description and relevant references is reported in Table 1.

**Table 1:** List of the strategies, named as the corresponding indices or algorithms, used to prioritize nodes for control optimization. See Fig. SI1 for a spatial representation.

| Strategy | Type | Description | References |
| --- | --- | --- | --- |
| In-degree | Network centrality index | The number of edges (connections) incoming a node | Pósfai and Barabási (2016) |
| Out-degree | Network centrality index | The number of edges outgoing a node | Pósfai and Barabási (2016) |
| In-strength | Network centrality index | The sum of the weights of the edges incoming a node | Pósfai and Barabási (2016) |
| Out-strength | Network centrality index | The sum of the weights of the edges outgoing a node | Pósfai and Barabási (2016) |
| Betweenness | Network centrality index | The number shortest paths between two other nodes of the network passing for a specific node | Freeman (1978) |
| Coreness | Network propagation index | The k-shell decomposition of a network into substructures from the most peripheral to the most central one | Seidman (1983) |
| GRWA | Network propagation index | Generalized Random Walker Accessibility: a measure of accessibility based on the Random Walker | Travençolo and Costa (2008); de Arruda et al. (2014) |
| VoteRank | Network propagation index | The ability of spreading information, based on an iterative voting algorithm | Zhang et al. (2016) |
| Vulnerability | Epidemic risk index | The average local losses by secondary infection, started anywhere in the region | Radici et al. (2024) |
| Dangerousness | Epidemic risk index | The overall average losses in the region caused by a primary infection in that node | Radici et al. (2024) |
| OrchardDen | Territorial index | The peach cultivated area within a node | - |
| Random | Stochastic algorithm | Random sampling | - |
| Regular | Stochastic algorithm | A sampling based on the `st_sample()` function (`type = 'regular'`) of the `sf` R package | - |
| VulnRand | Mixed stochastic algorithm | Stochastic sampling, weighted by vulnerability | - |
| DanRand | Mixed stochastic algorithm | Stochastic sampling, weighted by dangerousness | - |

Nodes indices are computed on the network defined by the matrix **W**.

### 2.3 Evaluating management performances

We evaluated the performance of the prioritization strategies for 126 intervals of treated nodes (ranging from 0 to 755 in increments of 6, *i.e.* 0, 6, 12, …750, 755) for a total of 14 × 126 = 1764 management scenarios. We define a management scenario (MS) as a combination of quantity of treated nodes prioritized with a given strategy. From an operational perspective, we: *i*) set the number of nodes to be treated; *ii*) set the prioritization index or algorithm and identify the corresponding nodes for treatment; *iii*) run the model to assess crop production based on contributions from both treated (*T*) and untreated (*U*) nodes. These three steps were repeated systematically to evaluate 1764 possible MSs. Due to the stochastic nature of the SEI model, we conducted 500 Monte Carlo simulations for each MS (for a total of 1764 × 500 = 882 · 10^3^ model runs) to gather robust statistics on crop production, including median values and percentiles. For each of the 500 replicates of a given MS, we randomized the peach variety distribution across the domain, the starting year (randomly selected between 2001 and 2010), and the initial infection state. Following Radici et al. (2024), the initial infection state assumes that 20% of the nodes are infected at the beginning of the simulation. For each MS, we assessed crop production *P_MS_*at harvest time in the 5^th^ simulated year, explicitly considering the contribute of treated and non treated nodes:

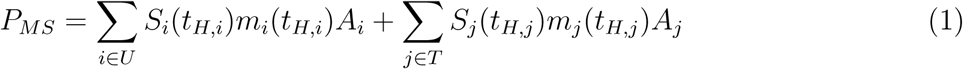

where *S* and *m* are respectively the susceptible fruit density (*fruit/m*^2^) and the single fruit mass (*g/fruit*, estimated via a a fruit growth curve from Bevacqua et al., 2023) at harvest time *t_H_*, whilst *A* is the peach cultivated area in treated (*i* ∈ *T*) or untreated (*j* ∈ *U*) nodes.

Peach cultivated areas varies between nodes, hence the same number of treated nodes may correspond to different treated areas. Meaningful comparison between strategies should be based on the total treated area. To robustly identify the prioritization strategy that performed best for a given level of fungicide application, we conducted two-sample Wilcoxon tests (Mann and Whitney, 1947) between groups of strategies for different treated areas (*i.e.*, under 10%, 20%, …, 100%). For each treatment threshold, we considered as optimal those strategies that never resulted in lower production, according to the Wilcoxon test.

Assuming that crop production from untreated areas may have higher market value than that from treated areas, we estimated a proxy of crop related revenues (*R_MS_*) for a given MS as:

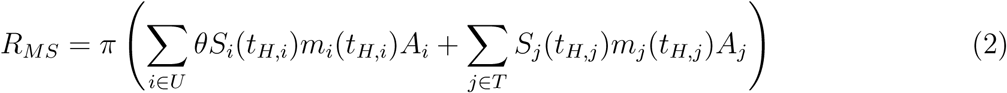

where *π* represents the price for peaches from treated areas and *θ* is a price multiplier for peaches from untreated areas (which we will refer to as “untreated fruit”). Since *π* is constant, we considered the proxy *R′_MS_* = *R_MS_/π* and analyzed the relative impact of different values of *θ* on *R′_MS_*. Specifically, we explored the threshold value of *θ* above which fungicide use becomes economically disadvantageous. The price multiplier indicates the consumer willingness to pay more for untreated fruit.

## 3 Results

National crop production *P_MS_* increases with the treated area (see Fig. 2), reducing disease spread, up to approximately 166 kton (16.7 t/ha), when all the peach cultivation areas (*i.e.*, all nodes in the network) are treated, which corresponds to a scenario with no disease. On the other hand, in absence of any treatment, the production is estimated to be around 133 kton, reflecting nearly 20% crop reduction. Our results allow for the identification of optimal prioritization strategy based on treated area. In Fig. 2a, the vulnerability index performs well for treated areas less than 20%, the out-strength index is most effective for treated areas between 25-35%, and the vulnRand index works best for treated areas between 45-85%. Beside being the most effective strategy for the largest share of treated area, vulnRand is also the best strategy overall (Fig. 2b). Stochasticity in the results decreases as the treated area increases, as it is primarily driven by the variability in epidemic dynamics, which is increasingly reduced as more nodes are treated (the interquartile rage decreases from 37 to 10 kton). For high treated areas (*>* 90%), it is not possible to identify a unique optimal strategy (see Table 2) because, as the number of treated nodes approaches the entire study area, the importance of the prioritization strategy diminishes. Interestingly, even in the 20-40% treated area range, no single strategy significantly outperforms the others, although both danRand and vulnRand perform well.

**Figure 2:**
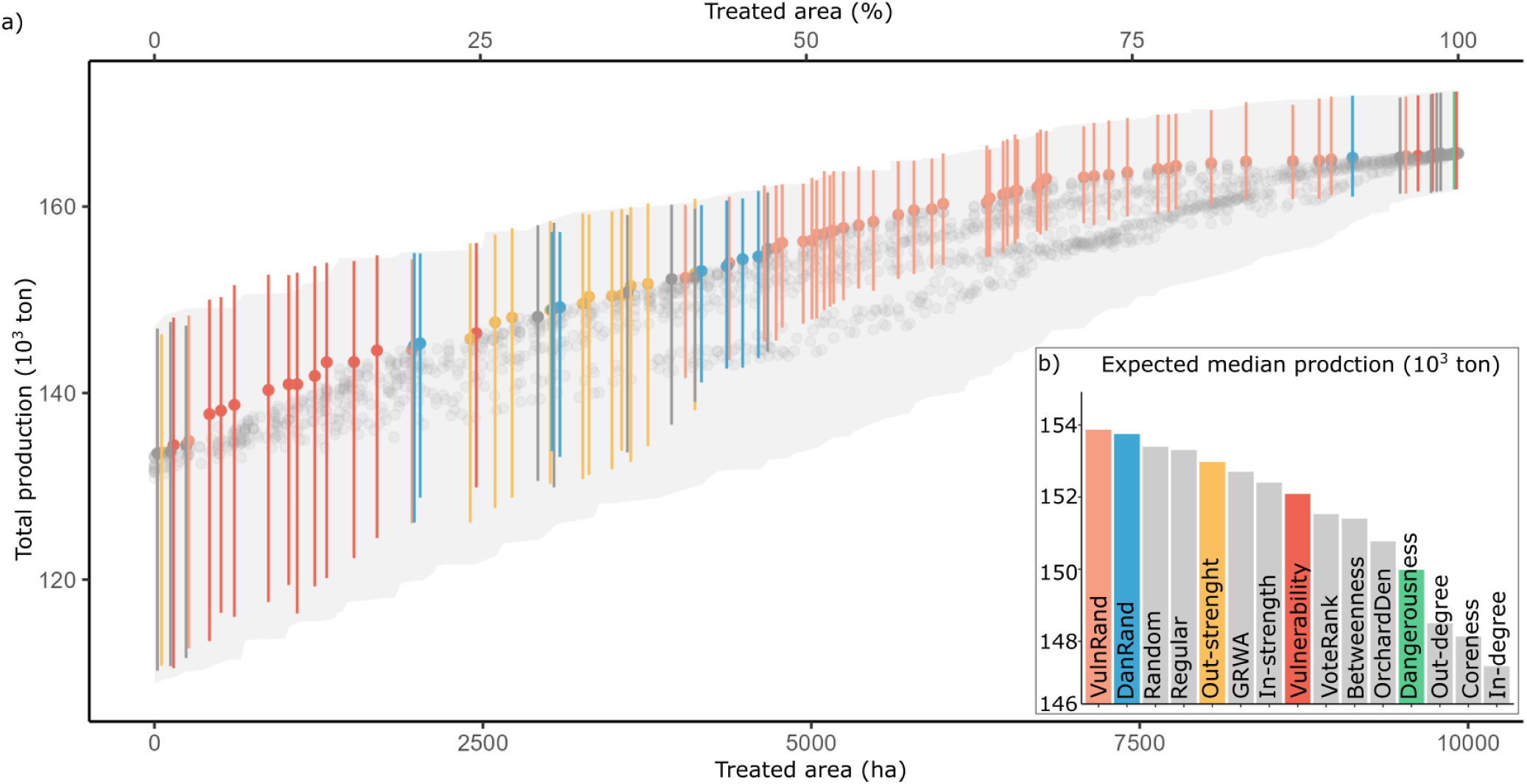
Estimated peach production *P_MS_* (median and inter-quartile ranges) for connected French cultivation areas, in the presence of brown rot disease, based on varying amounts of treated area where the disease cannot spread. The areas to be treated, modeled as nodes in a network, are selected according to different prioritization strategies (color code – see legend). The colored points represent the combinations of (treated area) × (prioritization strategy) that maximize total production, while the other combinations (median and inter-quartile ranges) are shown in grey. The inset (b) shows the expected value of the medians for each strategy (intended as the normalized integral over the trajectory of the medians) and provides a global view of each strategy’s performance.

**Table 2:** Statistically optimal strategies for each decile of treated area, *i.e.* whose performance are significantly better (p-value *<* 0.05 in the one-tail two-samples Wilcoxon statistical test) than any other. Two-sample comparisons are reported extensively in Fig.s SI2 and SI3.

| Treated area | Optimal strategies |
| --- | --- |
| 0 to 10% | Vulnerability |
| 10 to 20% | Vulnerability, orchardDen |
| 20 to 30% | DanRand, vulnRand |
| 30 to 40% | Out-strength, GRWA, danRand, vulnRand |
| 40 to 50% | VulnRand |
| 50 to 60% | VulnRand |
| 60 to 70% | VulnRand |
| 70 to 80% | VulnRand |
| 80 to 90% | VulnRand |
| 90 to 100% | DanRand, voteRank, vulnRand, Regular |

The curves of the maximum estimated revenues *R′_MS_*, relevant to national peach production, for different treated areas and price multiplier *θ* consider again a mix of prioritization algorithm (Fig.3). For *θ* = 1 (lower curve in panel a), there is no added value in untreated fruit, so that maximizing revenues is equivalent to maximize overall production (as reported in Fig.2) and could be obtained treating 100% of the production sites. For increasing values of *θ*, revenues increase, and the treated area at which the revenues are maximized (*i.e.* the treated area on the x-axis that corresponds to the highest point of the depicted curve) decreases until the extreme case, for *θ* = 2.9, it would be economically inconvenient to treat any single node. Since the estimated revenues vary with *θ*, also the optimal prioritization strategy for a given treated area varies. For *θ* = 1, the set of strategies optimizing production at a given treated area is the same as reported in Fig.2a. On the other hand, for increasing values of *θ*, strategies such as the out-strength, which performed well for *θ* = 1 for treated areas between 25-40%, are now outperformed. These outcomes are summarized in Fig.3b reporting the optimal treated area, together with the optimal prioritization strategy for node selection, as a function of *θ*. The strategies optimizing the total revenue are vulnRand (*θ* ∈ (1, 1.31]), danRand (*θ* ∈ (1.31, 1.73]), and vulnerability (*θ* ∈ (1.73, 2.88]). If *θ >* 2.88, the optimal management would be to leave all the crop production sites untreated.

**Figure 3:**
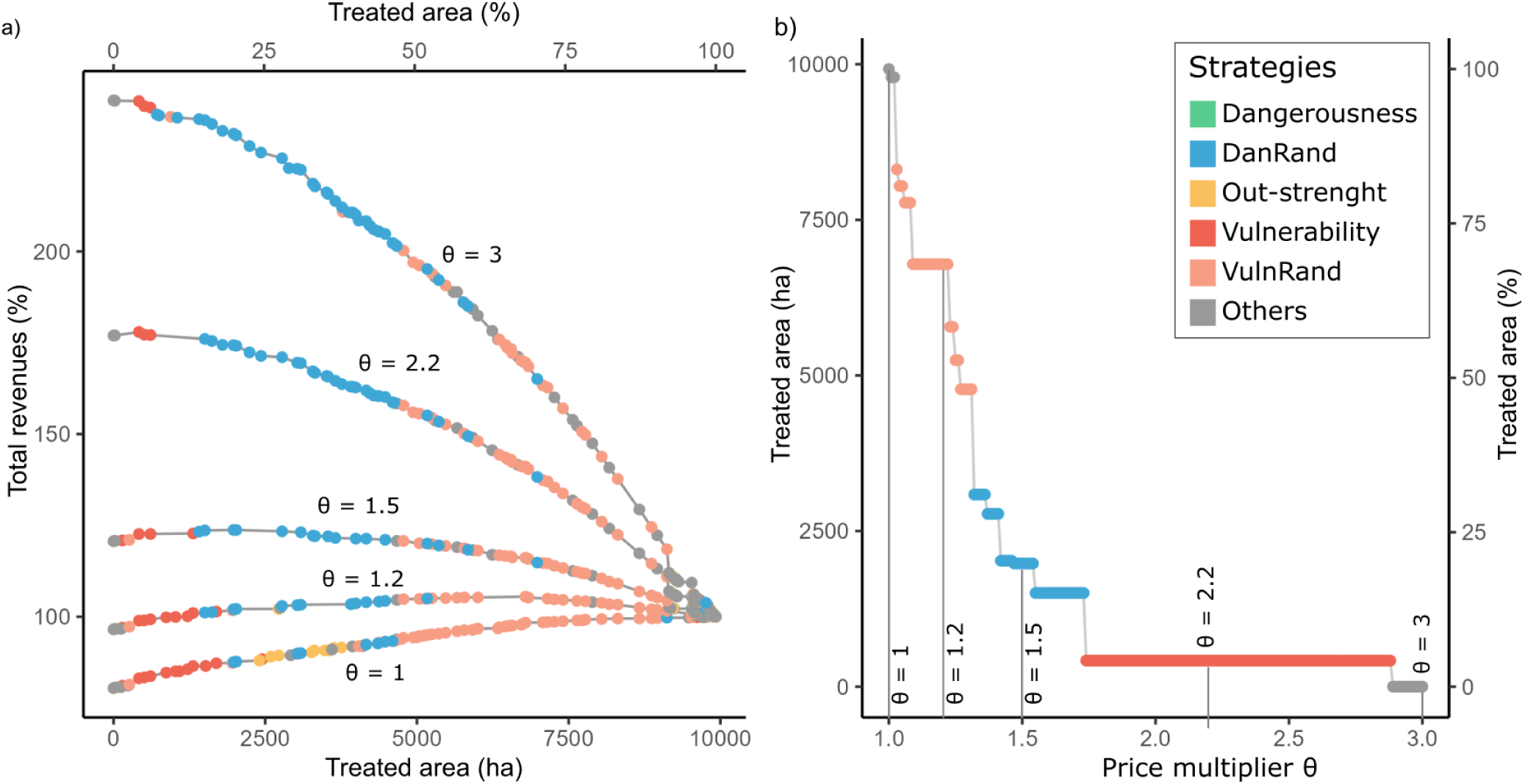
Effect of the variation of the price multiplier *θ* on the revenues and on the treated area. a) Estimated revenues *R′_MS_* variation for different treated areas and different market value of untreated fruit. The reference value is estimated via Eq. 2 for the extreme scenario where the entire peach production area is treated. b) Treated area and prioritization strategies maximizing revenues from peach production for different values of the price multiplier *θ*.

## 4 Discussion

This study is part of a body of recent research highlighting the importance of spatial planning at the landscape scale for effective disease management (Papäıx et al., 2014; Fabre et al., 2021). Dispersal of bioaggressors creates inter-dependencies between measures taken at different sites, necessitating coordinated strategies for optimized control. Landscape-scale management is essential for disease surveillance (Park et al., 2011; Carvajal-Yepes et al., 2019; Meyer et al., 2017; Radici et al., 2022) and it is increasingly recognized in EU agricultural policies (European Commission, 2020). For brown rot of peaches, we assumed that the national scale was appropriate for capturing these inter-dependencies. This choice represents a simplification, since bioaggressors spread over administrative boundaries. However, in this particular case, French boundaries corresponds to significative ecological obstacles (Mediterranean Sea, the Alps and the Pyrenees) to the dispersal of *Monilinia* spores. The main hypothesis would not have stand for other pathosystems, such as soybean rust (*Phakopsora pachyrhizi*) or wheat stem rust (*Puccinia graminis*), due to the large host crop areas and extreme pathogen mobility (Thompson et al., 2016; Radici et al., 2023b).

Increasing crop production and reducing the use of chemicals are considered conflicting objectives (Morris et al., 2024). However, in the long term, these objectives may converge due to the emergence of resistance against chemical treatments, which diminishes their effectiveness over time (Olitaa et al., 2023). Our model has provided insights into the form of the relationship between fungicide use and crop production. Specifically, the curve depicting optimized total production as a function of treated area and prioritization index exhibits a downwards concavity (Fig. 2a). This implies that in a scenario where all orchards are treated, a slight reduction in fungicide use would have a minimal impact on total production.

Strategy performances are characterized by very high stochasticity, implying the use of statistical tests to be robustly assessed. The first component of this stochasticity is related to the impact of weather and peach variety on total production. This is the only source of stochasticity when the domain is fully treated, and results in an interquartile range of about 10 kton in total production. The second component is epidemiological and concerns the probability of airborne infection, reflecting the randomness of wind as dispersal medium (Jeger et al., 2007). Combined with the first, the variability is highest when no treatment is applied (interquartile range of 37 kton).

Given the variety of available strategies to prioritize treatment sites, our findings can be conceptualized as identifying a Pareto front. This Pareto front highlights optimal management scenarios in which no alternative exists with both a smaller treated area and a higher total production. Among the prioritization strategies, the vulnRand index emerges as the most effective, thus composing large part of the Pareto front, particularly when the treated area exceeds 20%, with its random component gaining importance as treatment coverage increases. Presently, treated areas cover the vast majority of peach orchards (Ministère de l’Agriculture, 2020), but our analysis suggests that if agricultural practices were to shift toward a substantial reduction in fungicide use treating only a small fraction of sites — prioritization based on vulnerability would be the most effective. This case mirrors scenarios in public health where limited vaccine doses are preferentially allocated to the most vulnerable individuals. Similarly, in agriculture, prioritizing the treatment of the most vulnerable orchards under resource constraints serves to protect the areas at greatest risk. In the French context, this corresponds to targeting orchards in the middle Rhône region (Fig. 2 in Radici et al., 2024). A spatial node is considered more vulnerable if, while initially disease-free, it has a high risk of developing secondary infections (Meentemeyer et al., 2011). In our model, vulnerability is strongly influenced by site-specific factors, such as the frequency of rainy days during the susceptibility period, as precipitation is necessary for fruit to transition from an exposed to an infected state (Radici et al., 2024; Bevacqua et al., 2023).

Interestingly, the distribution of organic peach and nectarine cultivation across productive French departments partially aligns with our suggested strategy based on vulnerability (Ministère de l’Agriculture, 2020). For instance, in the Pyŕeńees-Orientales—a low-vulnerability region near the Spanish border (Fig. 2 and SI1)—44% of areas are organic. In contrast, the Drôme and Gard, which are highly vulnerable, have organic shares of just 5.9% and 6.1%, respectively. When the majority of the nodes in a network are treated, the strategy used to select which ones to treat becomes less significant, up to the extreme case where, if all nodes are treated, any ordering strategy becomes irrelevant. However, before reaching this extreme case, our results indicate that the composite index combining vulnerability with random sampling turns to be the best. In our study, random sampling alone performed unexpectedly well and incorporating a random component into an environmentally informed prioritization, such as vulnerability, enhanced performance further. Unlike spatially autocorrelated sampling techniques, random sampling achieves a more homogeneous node coverage, similar to regular sampling, which also shows a competitive performance (Fig. 2b). Random sampling is commonly used in disease surveillance to ensure broad coverage (Herrera et al., 2016), and combining different prioritization algorithms in a new one is a well-established practice in plant disease management (Sutrave et al., 2012). We propose that combining environmentally informed prioritizations (like vulnerability) with random sampling helps mitigate decentralized transmission of the disease, especially after high-risk nodes have been prioritized for treatment.

Among the network-based indices tested, only out-strength demonstrated optimal performance under intermediate treatment targets. This finding suggests that, for controlling brown rot in peaches, management strategies should prioritize epidemic or territorial characteristics—such as rainy-day frequency, which correlates with vulnerability—over purely network-based metrics. Despite the minor role of network-based indices in our study, not all performed equally. Out-strength and out-degree consistently outperformed in-strength and in-degree, respectively. This differentiation aligns with their epidemiological interpretations: nodes with high out-degree or out-strength are more likely to act as influential spreaders in small networks (Pautasso et al., 2010). Similarly, Andersen et al. (2019) demonstrated that the out-degree of a seed-trade network’s starting node can determine the final size of a disease outbreak. These observations underscore the utility of certain network metrics in identifying influential nodes, even if their broader application to our pathosystem is limited.

### 4.1 The rationale for higher prices of untreated fruit

Our exercise allowed us to identify which areas, in the hypothetical case of a collective management of the national peach production area, should remain untreated in order to maximize collective revenues. Nowadays, 88% of the French peach and nectarine cultivars are treated with fungicide (Ministère de l’Agriculture, 2020). According to our model, a value of *θ* ≈ 1.03 would justify such a shift. However, the price of peach coming from organic productions is almost double (*i.e. θ* ≈ 2; SelinaWamucii, 2023). One epidemiological reason for this mismatch is the fact that we modeled only the impact of brown rot, while fungicide-free peach orchards face risks from multiple fungal diseases, including peach leaf curl (*Taphrina deformans*), peach scab (*Cladosporium carpophilum*), and powdery mildew (*Podosphaera pannosa*) (Luo et al., 2022). Second, organic farming standards are far more rigorous than our modeled “untreated” category, and consequently, production could be more significantly reduced than what is expected in our model. In fact, organic farming not only addresses the use of fungicides but also regulates other phytosanitary products, the use of fertilizers, and other agricultural practices. Nonetheless, this does not diminish the importance of our analysis, since, its significance lies in demonstrating the effect that an higher market value of products from organic farming should have on the choices made in coordinated agricultural management. This value can be increased both by a greater willingness of consumers to pay and by higher costs associated with the use of polluting substances, according to the “polluters pay” principle. In fact, consumers are prone to pay more for products issued by low-input agriculture. Lin et al. (2008) estimated price premiums associated with product attributes focusing on five major fresh fruits and five major fresh vegetables in the United States and suggested that the organic attribute commands a significant price premium, which varies greatly from 20% above prices paid for conventional grapes to 42% for strawberries among fresh fruit and from 15% above prices paid for conventional carrots and tomatoes to 60% for potatoes. Similarly, de Souza Tavares et al. (2021) reported that organic juices are more expensive than their conventional counterparts with prices approximately 50% higher in Brazil and 10% higher in France, supporting the perception of higher costs associated with organic products. Alternatively, higher income to producers who favor untreated production might be induced by policies designed to internalize the environmental costs of synthetic compounds used in treatments (Kümmerer et al., 2019). According to the “polluters pay” principle, external costs could be internalized through taxation, promoting less harmful alternatives (Ambec and Ehlers, 2016). This principle underpins policies like the “carbon tax”, which aims to reduce greenhouse gas emissions by increasing the costs of high-emission options (Khan, 2015).

Although the higher profits associated with organic fruit production may encourage the expansion of these practices, especially in the wealthiest parts of the world, it is important to remember that bioaggressors control aims to ensure food access for all of the planet’s inhabitants (Savary et al., 2019). This means that, when an agricultural product is essential for feeding a population, it is unacceptable to reduce its quantity in order to increase producers’ profits. While political considerations are beyond the scope of our work, we believe they should not be overlooked when interpreting our results.

Apart from the omission of other peach fungal pathogens, an important limitation of the study concerns the absence of fungicide-related costs. These costs, if taken into account, would make the production of untreated fruit easier than in the presented framework. This would result in a faster reduction of the treated area as the price multiplier *θ* increases.

When considering the impact of bioaggressors in crop production in future decades, particular attention should be given to the synergy with expected climate change, which is already affecting agricultural yields and larger impacts are expected in future scenarios (Ray et al., 2019; Ortiz-Bobea et al., 2021; Lesk et al., 2016). Estimates indicate an expected reduction of the global crop yield from 3% to 7% for each degree-Celsius of temperature increase (Zhao et al., 2017; Liu et al., 2016; Rezaei et al., 2023) and a reduced cropland suitability at lower latitudes (Rosenzweig et al., 2014; Zabel et al., 2014). In the case of peach production in France, it is expected to decline drastically without adaptation measures, as regions where peach is currently cultivated might not be able to assure chilling conditions necessary for plant blooming (Vanalli et al., 2021). Conversely, the impact of brown rot may diminish under warmer and drier conditions (Vanalli et al., 2024).

## Acknowledgement

The authors acknowledge the support of funding from the French National Research Agency (ANR) for the BEYOND project (contract # 20-PCPA-0002) and the SuMCrop Sustainable Management of Crop Health Program of INRAE that supported the work of all authors.

We thank Nik Cunniffe, Chiara Vanalli and Ghislain Génieux for constructive comments and the colleagues of the AgroClim INRAE unit who developed the SICLIMA platform, which allows to access climatic reanalysis as computed by Météo-France.

## Data Availability Statement

The code of the model supporting the study is available at https://github.com/radiciandrea/MetapopulationBrownRot.

## Declaration of generative AI and AI-assisted technologies in the writing process

During the preparation of this work, the authors used ChatGPT-4 and DeepL in order to improve the readability and language of the manuscript. After using this tool/service, the authors reviewed and edited the content as needed and take full responsibility for the content of the published article.

## Supplementary information

### 1 Prioritization strategies

In Fig. 1 we summarize the values of each index for each prioritization strategy - with the exception of voteRank, which provides directly the ranking itself. Random, regular, vulnRand and danRand are not shown since they are based on stochastic algorithms, and can not be mapped consequently. In particular, random, vulnRand and dandRand are based on the `sample` function of the `base` library of `R`, while regular is based on the `st_sample()` function (with `type = ‘‘regular’’`) of the sf package.

### 2 Statistical tests

In Fig.s 2 and 3, we present the results of the two-samples Wilcoxon tests (Mann and Whitney, 1947) among groups of alternatives divided by decile of treated area. Optimal alternatives are summarized in Tab. 2 in the main text.

---

**Figure 1:**
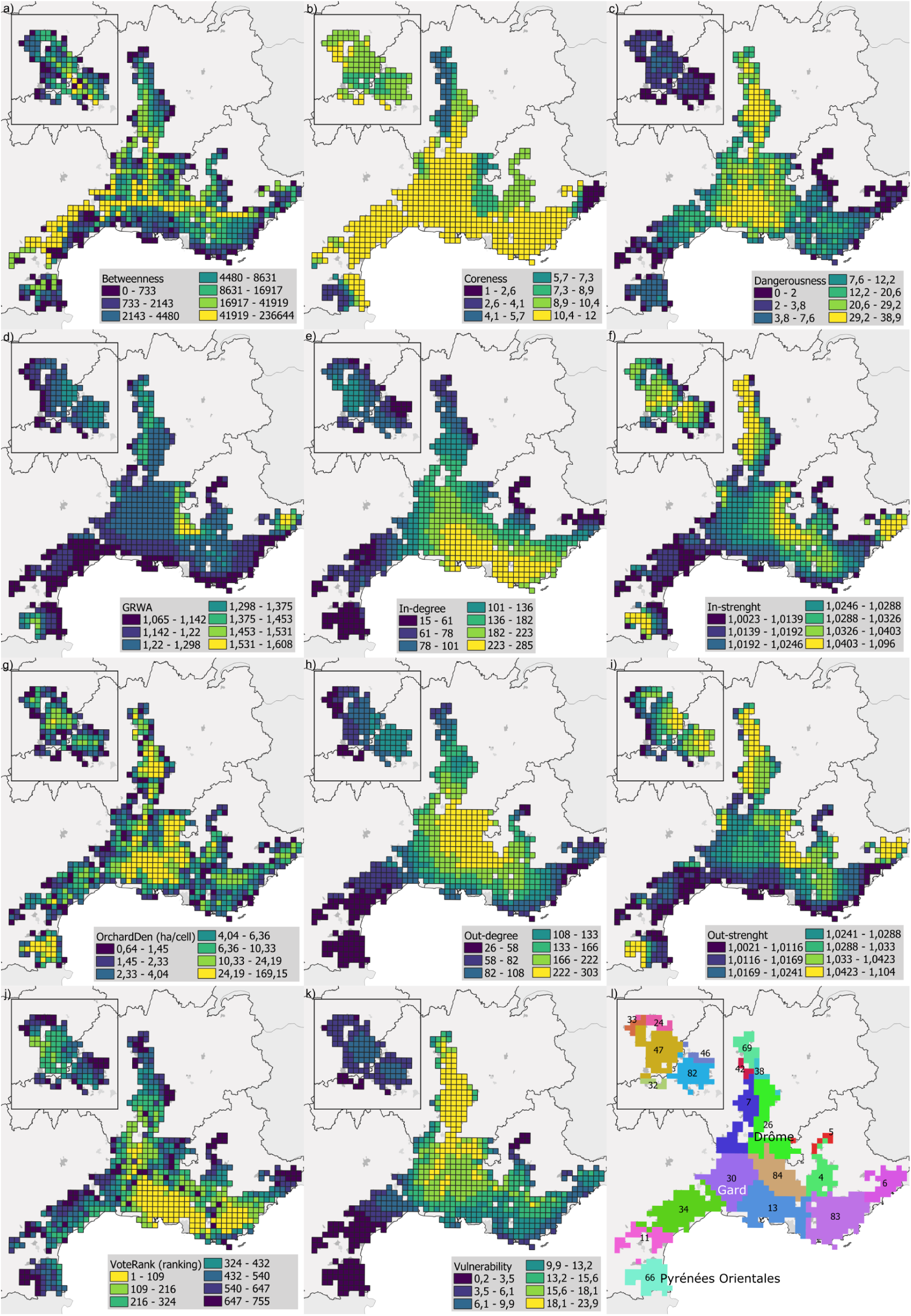
Value of prioritization indices (for voteRank, panel j, we show the value of the ranking itself). In panel l, we show a map with the departments.

**Figure 2:**
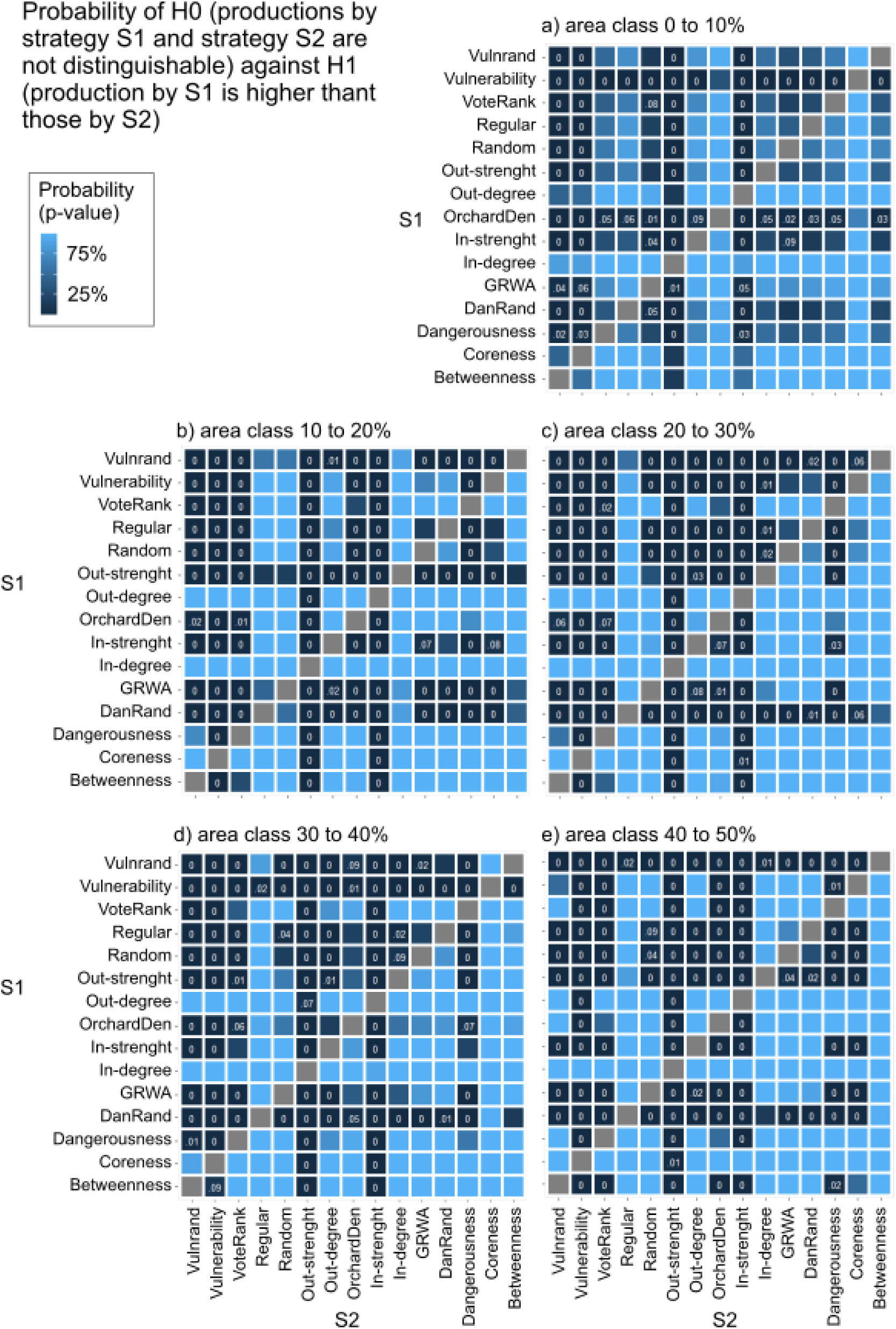
Results of the statistical two-samples Wilcoxon tests, area classes 0 to 50%. To improve readability, only p-values *<* 0.05 are shown in numbers.

**Figure 3:**
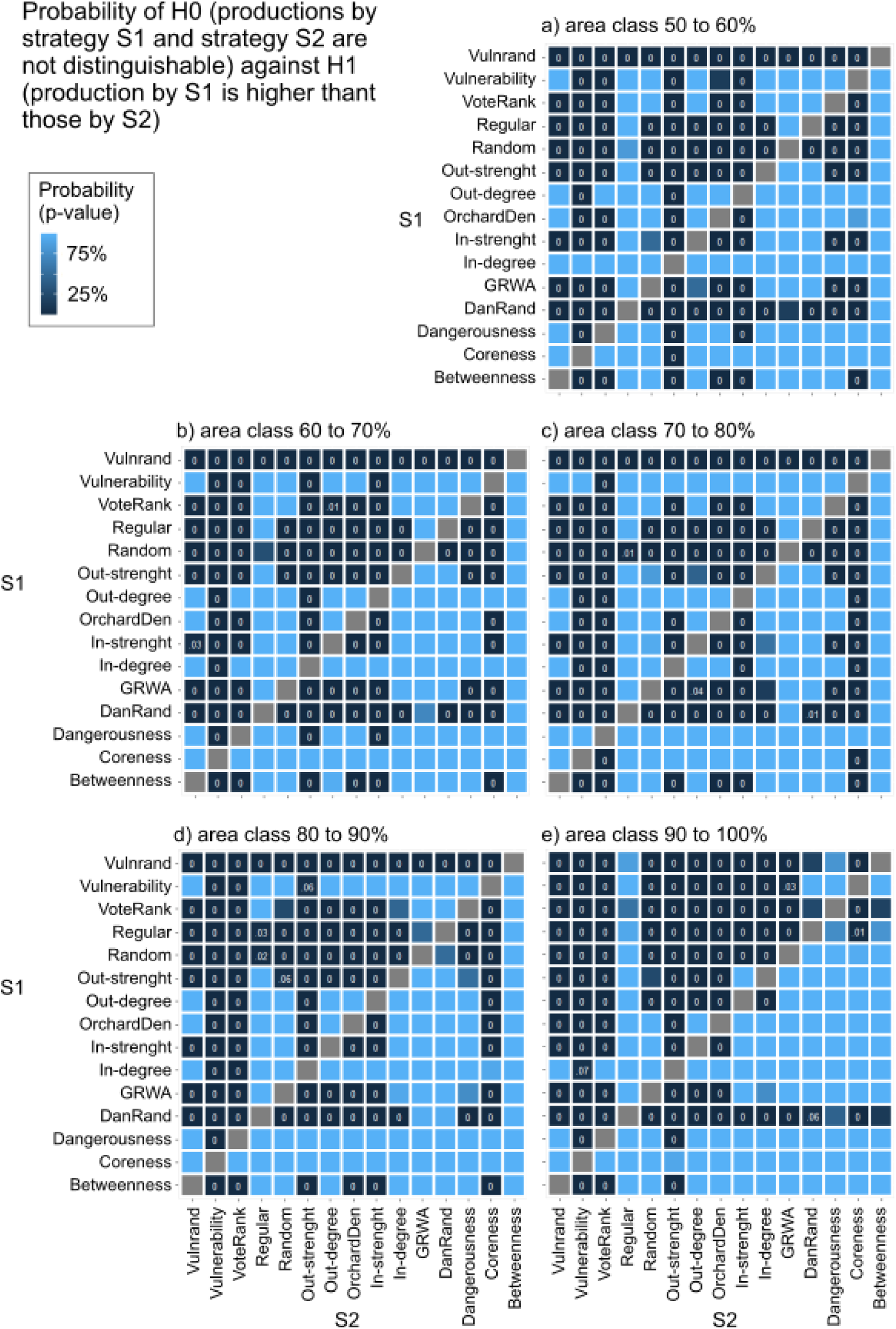
Results of the statistical two-samples Wilcoxon tests, area classes 50 to 100%. To improve readability, only p-values *<* 0.05 are shown in numbers.

